# Detect *de novo* expressed ORFs in transcriptomes with DESwoMAN

**DOI:** 10.1101/2025.06.10.658796

**Authors:** Anna Grandchamp, Marie Lebherz, Elias Dohmen

## Abstract

*De novo* gene emergence refers to the process by which new genes arise from mutations in previously non-coding genomic regions. Prior to becoming fixed in a species, newly expressed open reading frames (neORFs) undergo significant turnover within their species of origin. Investigating these early stages of *de novo* gene emergence is essential for understanding the mechanisms that enable gene formation from scratch. No software currently exists that can identify and characterise novel, unannotated open reading frames from a transcriptome, and analyse their mutations and fixation patterns within or across species.

To address this gap, we introduce **DESwoMAN** (***D****e novo* **E**mergence **S**tudy **W**ith **O**utgroup **M**ut**A**tio**N**s), a software tool designed to: (1) detect neORFs in transcriptomes, (2) filter neORFs with no homology to outgroup genes, and (3) search for syntenic sequences homologous to neORFs in outgroup genomes (and optionally transcriptomes) and analyse mutations in coding features between these sequences. We applied **DESwoMAN** with two different strategies to three setups, using twice human and once fruit fly as query species. Our results highlight the tool’s capabilities and demonstrate its potential for elucidating the early stages of *de novo* gene emergence.

**DESwoMAN** is available at https://github.com/AnnaGrBio/DESWOMAN. It is implemented in Python3 and comes with a docker image on DockerHub for easy installation and execution including all (non-Python) dependencies.

## Introduction

*De novo* gene birth is the process by which a non-genic region acquires genic features by mutation (Zheng and Zhao, 2022; Zhao et al., 2014; Zhao, 2023; Van Oss and Carvunis, 2019; Vakirlis et al., 2020; Rich and Carvunis, 2023; Parikh et al., 2022; Vakirlis et al., 2022; Wissler et al., 2013; Schmitz et al., 2018). According to the model proposed by Carvunis et al. (2012), the emergence of genes from scratch follows two main steps: First, a genome acquires by mutation an open reading frame (ORF) and transcription. Second, this transcribed ORF becomes fixed at the species level. If such a transcribed ORF is translated, but is not fixed in the species, the new gene is qualified as a proto-gene. The proto-gene’s fixation stage is likely very dynamic, as several studies have demonstrated a high turnover in gain and loss of recently gained new transcripts and ORFs within a species (Grandchamp et al., 2024; Iyengar and Bornberg-Bauer, 2023). However, proto-genes can become subject to selection pressure (Li et al., 2010; Palmieri et al., 2014; Wacholder et al., 2023; Ward and Kellis, 2012) and some of them become therefore fixed in a species. Such genes are called *de novo* genes and can be detected in species or phylogenetically restricted groups (Peng and Zhao, 2024; Vakirlis et al., 2018; Vakirlis and McLysaght, 2019; Weisman, 2022).

In this paper, we call newly expressed ORF (neORF) a proto-gene that was detected in silico, for which there is no evidence of translation. Depending on the region of emergence, different mutations may be required for the birth of an neORF. These mutations can be the emergence of the ORF by mutations leading to a start or stop codon for example, the emergence of transcription initiation sites, a combination of nucleotides that provide stability to untranslated regions (UTRs) and allow translation or introduce splicing sites in the case of introns. Validating a *de novo* gene emergence and understanding the underlying mechanisms remains a methodological challenge as their initial mutations are difficult to determine. However, these mutations are crucial to study the function and properties of genes arising through this mechanism. To infer the *de novo* emergence status of annotated genes in a genome, several bioinformatic tools have been developed (Heames et al., 2020; Vakirlis et al., 2018; Wu and Knudson, 2018; Wang et al., 2020; Zhuang and Cheng, 2021; Wu et al., 2011; Prabh and Rödelsperger, 2019; Murphy and McLysaght, 2012; Yang and Huang, 2011; Knowles and McLysaght, 2009; Neme and Tautz, 2013; Moyers and Zhang, 2016; Cai et al., 2008; Peng and Zhao, 2024). In 2019, Vakirlis and McLysaght (2019) developed protocols to validate the *de novo* emergence of annotated genes in genomes and implemented filtering steps such as removing candidates with coding homologs not annotated in outgroup genomes and reconstructing the ancestral state of the *de novo* candidate. The software DENSE (Roginski et al., 2024b) uses annotated genes and the corresponding genome as input, validates the lack of detectable homology to any known protein in the NCBI NR database (Sayers et al., 2019), and searches for homologous non-genic hits in outgroup genomes. All these pipelines work with annotated *de novo* genes, but the earliest stages of such genes are typically missed by traditional gene annotations. More precisely, genome annotation pipelines (Gabriel et al., 2023) use gene homology or known genic features to annotate genes in new genomes. However, neORFs neither have detectable homology to other genes, nor exhibit known genic features and are therefore missed by such an approach.

To detect early stages of genes and validate their *de novo* emergence status, several studies have used transcriptomes to search for neORFs as *de novo* gene candidates (Dowling et al., 2020; Schmitz et al., 2018, 2020; Ruiz-Orera et al., 2015; Blevins et al., 2021; Sandmann et al., 2023; Zhang et al., 2019; Zhao et al., 2014; Vakirlis et al., 2022; Witt et al., 2019; Grandchamp et al., 2024, 2023; Zhao, 2023; Schmitz et al., 2018; Neme and Tautz, 2016). However, the specific methodology differs significantly between studies (Dohmen et al., 2025) and it partly requires high computational skills and extensive decision-making at each step of this long process (Grandchamp et al., 2025) to reproduce the resulting annotations or analyse other input data the same way. Furthermore, the majority of these approaches do not investigate the mutations that lead from a non-coding state to an neORF. This step can be achieved through the extraction and comparison of syntenic non-genic homologs in closely-related outgroup genomes.

Detection and validation of the earliest stages of *de novo* gene emergence is the first and most important step to understand the mechanisms underlying gene birth from scratch. Knowledge of these molecular mechanisms will help us to better understand the evolution of completely novel functions and the beginning of life. Here, we present **DESwoMAN** (***D****e novo* **E**mergence **S**tudy **W**ith **O**utgroup **M**ut**A**tio**N**s), a standardised and fully automated pipeline designed to automatically detect neORFs based on transcriptomes, validate their *de novo* status, and extract syntenic homologous regions to neORFs from outgroup genomes. Based on the extracted syntenic homologous sequences, **DESwoMAN** identifies different mutations responsible for the coding or non-coding status of a sequence within the same species or between closely related species.

## Results

### Identifying neORFs with DESwoMAN

We developed **DESwoMAN**, a software to detect, validate and analyse properties of newly expressed ORFs (neORFs). In order to ascertain the practical benefits of the software and to gain novel insights about *de novo* emergence mechanisms, we apply **DESwoMAN** in this study to three different setups with two different strategies.

Strategy 1 allows to detect neORFs in a transcriptome, and study the mutations that could have led to its emergence by searching syntenic homologues in outgroup genomes. The HumanSetupS1 serves as a dataset to study neORFs in *Homo sapiens* in comparison to other mammalian species, while the DrosoSetupS1 serves as a dataset to study neORFs in *Drosophila melanogaster* at the population level.

HumanSetupS2 serves as a dataset for Strategy 2 that allows to detect neORFs in several transcriptomes from one single species, and study common expression.

### Detecting mutations in neORFS with Strategy 1

We detect human-specific neORFs (HumanSetupS1) and fruitfly population-specific neORFs (DrosoSetupS1) by applying DESwoMAN’s Strategy 1. We find 1,562 human-specific intergenic neORFs and 898 specific intergenic neORFs from a transcriptome of the Finland population (FI) of *D. melanogaster*. For 63.44% to 71.76% of the 898 *D. melanogaster* ‘s neORFs, DESwoMAN could detect syntenic homologous sequences in the other genomes of *D. melanogaster* (Table 1). In human we find for 68.37% to 92.2% of the neORFs syntenic homologous sequences in primates. In mice, however, only for 1.0% of the human neORFs syntenic homologous sequences are detected. Contrary to our initial assumptions, we detect more syntenic homologous sequences for human neORFs in other primate species than we detect for *D. melanogaster* neORFs in other populations of the same species.

**Table 1:** Number of neORFs and their syntenic homologous sequences.

| sample/species | # syntenic homologs | % syntenic homologs |
| --- | --- | --- |
| Query : FI | 898 | - |
| DK | 991 | 63.44 |
| ES | 1121 | 71.76 |
| SE | 1060 | 67.86 |
| UA | 1069 | 68.43 |
| TR | 1038 | 66.45 |
| ZI | 1013 | 64.85 |
| Query : human | 1562 | - |
| Chimpanzee | 827 | 92.09 |
| Bonobo | 828 | 92.20 |
| Gorilla | 819 | 91.20 |
| Orangutan | 757 | 84.29 |
| Macaque | 614 | 68.37 |
| Mouse | 9 | 1.00 |

The conservation of coding features shows little variation in syntenic homologous sequences across all *D. melanogaster* samples (Fig. 1), while it is more variable between mammalian species in the HumanSetupS1. As expected, coding features of syntenic homologous sequences exhibit reduced conservation with increased phylogenetic distance. Start and stop codons are, on average, fairly well conserved within syntenic homologous sequences (https://zenodo.org/records/14936107) with an exception in the far-related mouse. Contrary to our expectations, start and stop codons exhibit a greater degree of conservation within species that are closely related to human than between populations of *D. melanogaster*.

**Figure 1:**
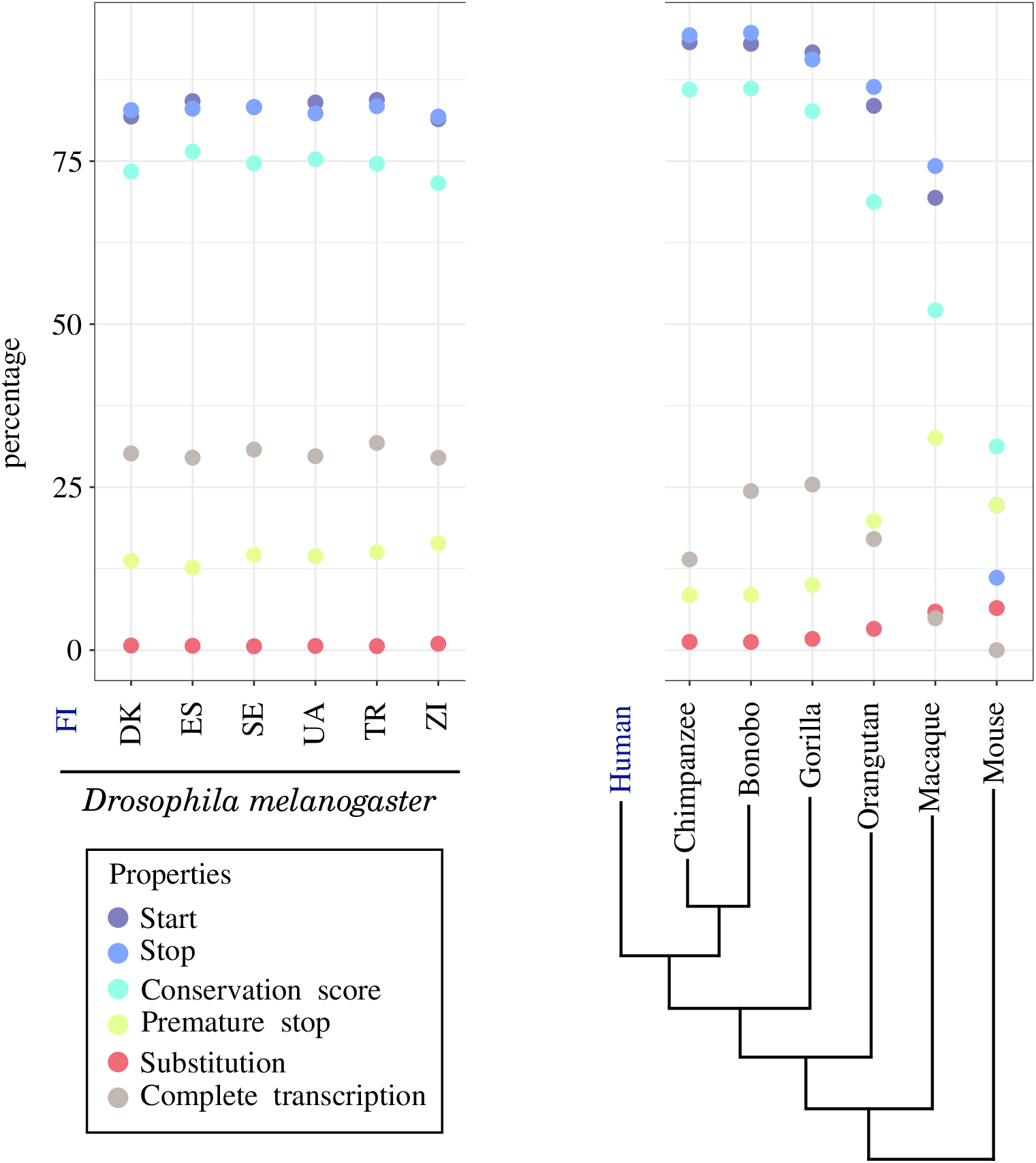
Conservation of coding features of syntenic homologous sequences as percentage of the total number of neORFs that exhibit: a start codon, a stop codon, a premature stop codon, or complete transcription. Conservation score represents the average frameshift conservation score of all syntenic homologous sequences of neORFs. Substitution represents the average percentage of substitutions in syntenic homologous sequences in comparison to their respective neORFs across all neORFs.

Transcription, on the other hand, is more conserved between *D. melanogaster* populations, ranging from 29.52% to 31.79% of syntenic homologous sequences being fully transcribed in the same orientation. The transcription status of syntenic sequences homologous to human neORFs is lower in mammalian species, ranging from 4.88% to 25.39%, excluding mouse. Conservation scores based on frameshift were high on average in all fruit fly (71.64% to 76.48%) and range from high to low in outgroup species of humans with increasing phylogenetic distance (86.14% to 52.12% excluding mice). The average percentage of substituted nucleotides in syntenic homologous sequences of neORFs is always very low.

**DESwoMAN** applies a synteny window to detect syntenic homologous sequences of intergenic neORFs, which can be adjusted/configured by the user. To determine synteny, homologous conserved genes are used as anchors and the window size specifies how many of these anchors up-or downstream of the neORF are considered. Moreover, syntenic homologs can be detected by using simple reciprocal BLAST hits, or by using best reciprocal BLAST hits. Simple reciprocal BLAST hits refer to any two sequences that are found through a BLAST search, while best reciprocal hits are a stricter subset with only the top-scoring sequence of the respective BLAST search.

We investigate whether the size and flexibility of the synteny window for intergenic neORFs affects the number of syntenic homologous sequences that are detected (Fig. 2, https://zenodo.org/records/14936107). We test 5 different synteny window sizes on the two setups, as well as a no-synteny option. With simple reciprocal hits (Fig. 2 a and c), we detect more syntenic homologous sequences of human neORFs than of fruit fly neORFs in outgroup genomes. In both setups we observe that the larger the synteny window is, the higher is the number of syntenic homologous sequences. In fruit fly we find on average 37% of neORFs in other fruit fly samples with a synteny window of 1, which increases to an average of 71% with a synteny window of 5 (Fig. 2 a). In the human setup, neORFs have on average 55% of syntenic homologous sequences in outgroup species (excluding mice) with a synteny window of 1, and this increases to 87% on average with a synteny window of 5 (Fig. 2 c, https://zenodo.org/records/14936107).

**Figure 2:**
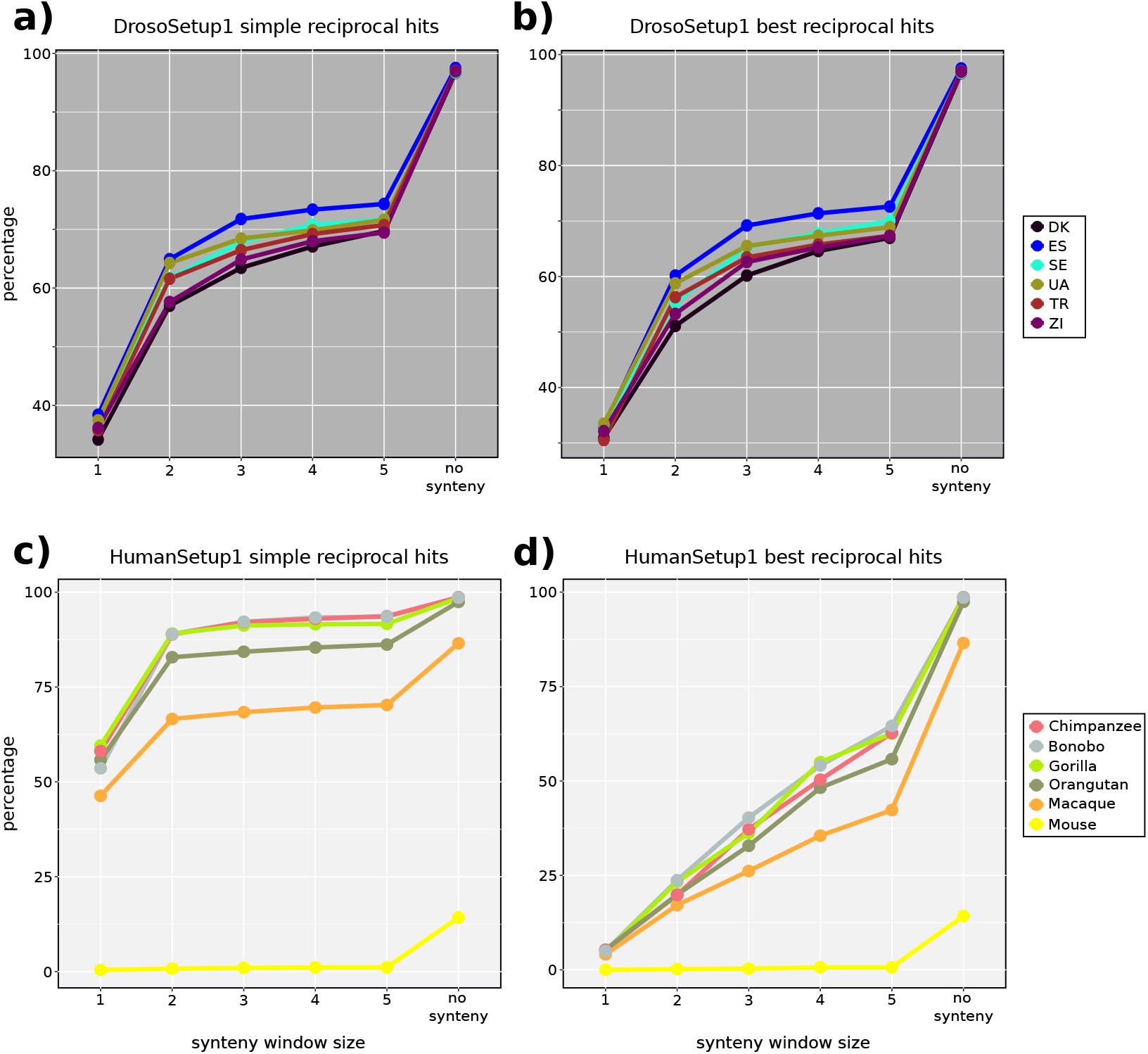
Impact of synteny window size and homology validation. (a) DrosoSetupS1 with simple reciprocal hits, (b) DrosoSetupS1 with best reciprocal hits, (c) HumanSetupS1 with simple reciprocal hits, (d) HumanSetupS1 with best reciprocal hits.

For both HumanSetupS1 and DrosoSetupS1, the absence of synteny criteria resulted in nearly 100% (96% to 97% for fruit flies and 86% to 98% for human outgroups) of detected syntenic homologous sequences, except in mice where 14% of neORFs are found as syntenic homologous sequences in the genome. In the HumanSetupS1 we observe that the greater the phylogenetic distance is, the fewer syntenic homolgous sequences were detected for all synteny windows (Fig. 2 c, https://zenodo.org/records/14936107).

With best reciprocal hits (Fig. 2 b and d), as with simple reciprocal hits, we observe that the larger the synteny window is, the more syntenic hits are detected. However, while the percentage of syntenic homologous hits remains almost unchanged for all fruit fly outgroups for any synteny window https://zenodo.org/records/14936107), a much more pronounced decrease was observed for human outgroups. For example, compared to simple reciprocal hits, the number of homologous hits observed for a window size of 2 with simple reciprocal hits shows an average of 83% between outgroup species, excluding mice, whereas this average percentage of syntenic homologous hits drops to 21% with best reciprocal hits. In the HumanSetupS1, the phylogenetic distance influences the amount of homologous hits in the same way.

### Analysis of tissue-specific neORF expression with Strategy 2

We applied **DESwoMAN**’s Strategy 2 to HumanSetupS2 to detect neORFs and assess how their transcription differs across tissues. Three independent groups of neORFs are investigated: intergenic, intronic, and antisense neORFs.

Across all the studied tissues, the same pattern of neORF frequency has been identified: antisense neORFs are most numerous, followed by intergenic ones, and intronic ones are the least numerous (Table 2). We find most neORFs across all three categories in cerebellum and fewest neORFs in testis.

**Table 2:** neORFS per transcriptomes.

| Transcriptome | total transcripts | genomic positions | # transcripts | ORFs | neORFs |
| --- | --- | --- | --- | --- | --- |
| **Human brain** | 32851 | antisense | 5920 | 11677 | 3931 |
|  |  | intergenic | 1483 | 2080 | 905 |
|  |  | intronic | 488 | 635 | 299 |
| **Human cerebellum** | 48810 | antisense | 9774 | 16216 | 6183 |
|  |  | intergenic | 4479 | 5940 | 2766 |
|  |  | intronic | 2272 | 3013 | 1326 |
| **Human heart** | 25313 | antisense | 4699 | 7658 | 2991 |
|  |  | intergenic | 539 | 615 | 311 |
|  |  | intronic | 442 | 432 | 231 |
| **Human kidney** | 32664 | antisense | 5932 | 8595 | 3629 |
|  |  | intergenic | 969 | 1170 | 533 |
|  |  | intronic | 448 | 461 | 241 |
| **Human testis** | 18231 | antisense | 2431 | 3817 | 1593 |
|  |  | intergenic | 288 | 326 | 173 |
|  |  | intronic | 90 | 82 | 46 |

All identified neORFs are classified into orthogroups by **DESwoMAN** (Fig 3). Across all three neORF categories (antisense, intergenic and intronic), the vast majority of neORFs is tissue-specific, with a consistent decline in the number of orthogroups shared by a higher number of tissues.

**Figure 3:**
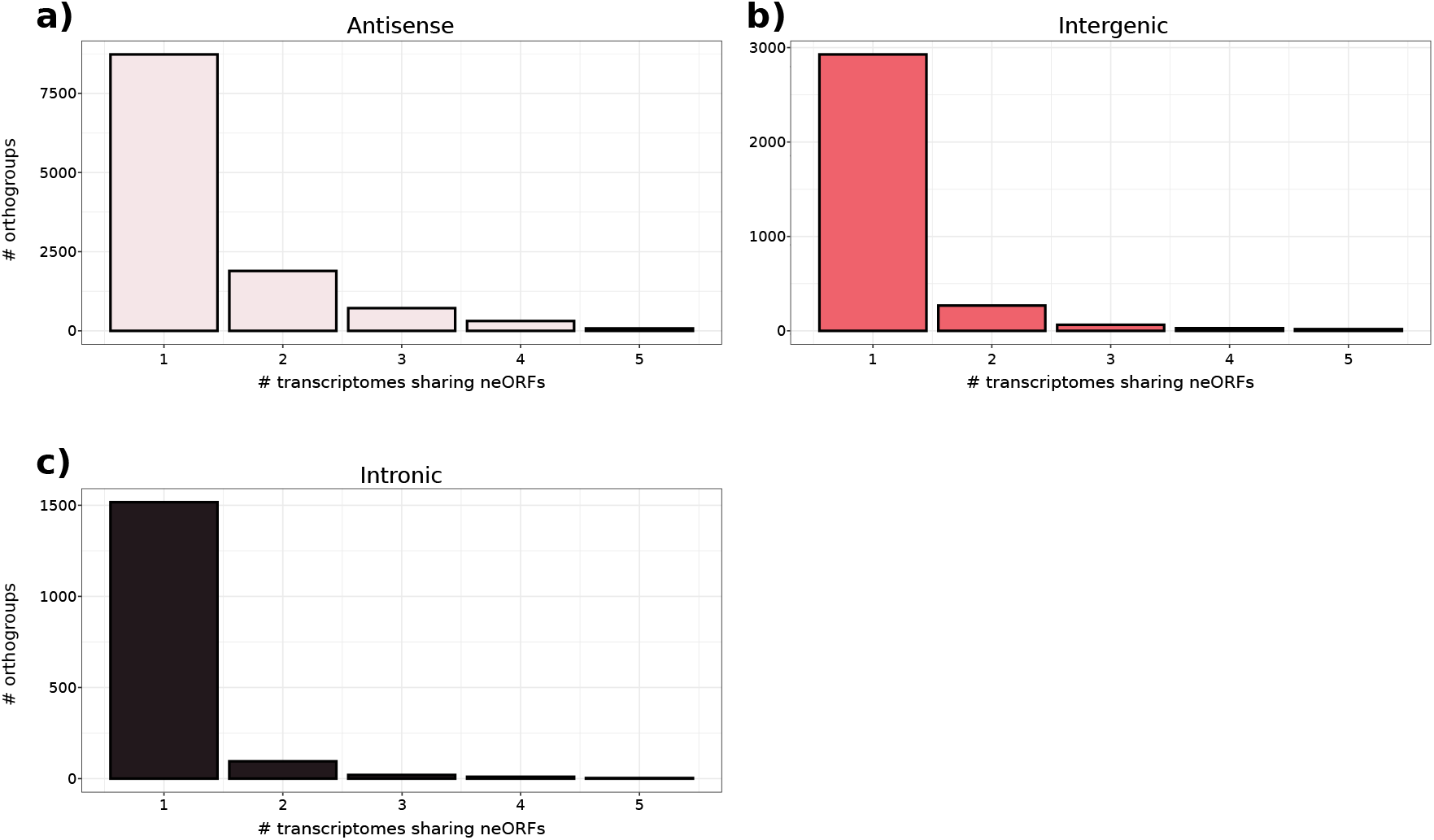
Tissue-specificity of neORF orthogroups in human.

## Discussion

### DESwoMAN: Automated neORF detection and analysis

In this study we present **DESwoMAN**, a tool developed to automatise the detection and analysis of newly expressed open reading frames (neORFs) based on transcriptome data. Our tool offers a high flexibility regarding the addressable biological questions by providing two different strategies with user-adjustable parameters to cover a wide range of use cases. In the three different use cases presented in this study, we gain biological insights about the earliest stages of *de novo* gene emergence through inter-species comparisons, as well as intra-species comparisons at the population level and regarding tissue-specificity of neORFs.

**DESwoMAN** reports multiple downstream analyses results next to the identified neORFs, such as syntenic homologous sequences and properties of the neORFs and their syntenic homologous sequences in the form of mutations associated with the coding status. However, the biological interpretation of the reported results remains a crucial task to be carried out by the user. For example, if all syntenic homologous sequences of a neORF reported by **DESwoMAN** share the same coding status, there is a high likelihood that the neORF does not represent a *de novo* gene, but rather a conserved gene, which is not yet annotated.

It is important to note in this context that the definition, detection and validation of a potential *de novo* status is already highly variable across studies ((Vakirlis et al., 2020; Vakirlis and McLysaght, 2019; Roginski et al., 2024a; Parikh et al., 2022)). **DESwoMAN** validates neORFs by identifying syntenic non-coding homologous sequences, which serve as a baseline for confirming a *de novo* emergence in most of the studies in the field.

Furthermore, many other different metrics can be used to assess coding status conservation, such as ancestral reconstructions (Vakirlis et al., 2024) or phylogenetic tools combined with protein sequence homology detection (Sandmann et al., 2023). **DESwoMAN** analyses 6 features to validate the coding status based on sequence alignments. Other potentially relevant features, such as splicing site conservation or translation status, are not determined by our tool. However, **DESwoMAN** reports genomic coordinates of syntenic homologs, allowing users to extract neORFs and their syntenic homologous sequences from genomes to apply alternative conservation metrics.

Apart from varying definitions and validation of a potential *de novo* and coding status, e.g. (Vakirlis et al., 2020; Vakirlis and McLysaght, 2019; Roginski et al., 2024a; Parikh et al., 2022)), several factors can influence the reliability of the results. If a low number of target genomes is used, for example, a missing coding status in all syntenic homologous sequences of an neORF could be misinterpreted, if a syntenic homologous coding ORF exists in other genomes, not included in the study. A higher number of well-chosen query and target genomes or transcriptomes can therefore lead to more robust conclusions.

### Strategy 1: neORF properties across species and populations

In order to demonstrate the functionality of **DESwoMAN**, we applied Strategy 1 to the HumanSetupS1 for comparison across species and to the DrosoSetupS1 for comparison across populations within a species.

Syntenic homologous sequences of neORFs are on average well conserved across all tested species or populations. They exhibit low substitution rates when compared to their corresponding neORFs, but lack in most cases at least one coding feature. The most common missing feature is transcription, with more than 70% of syntenic homologous sequences being not or not fully transcribed. This pattern is observed in both Strategy 1 setups and supports a high turnover in transcription gains and losses (Grandchamp et al., 2024; Clark et al., 2011). Furthermore, this finding underscores the role of transcriptional activation in the early stages of gene emergence (Neme and Tautz, 2016), even though some of these cases might be false negatives *de novo* gene could have low expression levels.

Another general pattern we can observe is a lower number of conserved coding features in syntenic homologous sequences with a greater phylogenetic distance to the query species, suggesting a low fixation rate of coding features. Interestingly, start and stop codons are not conserved 10 to 20% of the cases, and conservation scores are around 80%, while substitutions are often really low in syntenic homologs (1% of the sequences). These findings could indicate a higher mutation rate on coding features and important force of selection (Back, 1994), supporting the findings of previous studies (Lebherz et al., 2024b; Zhao et al., 2024; Schlötterer, 2015).

### Impact of synteny window size

We investigate the impact of synteny window size and BLAST hit detection regarding the accuracy at detecting syntenic homolog sequences of intergenic neORFs. Across all tested setups, larger synteny windows result in the detection of more syntenic homologous sequences. Several evolutionary mechanisms can make a larger window size necessary to find the surrounding, homologous genes, such as genome reshuffling, inversions, duplications, gene losses, horizontal gene transfer, and others (Zhang et al., 2023; Steenwyk and King, 2024). However, there is a trade-off between window size and accuracy. Increasing the window size can increase the number of identified sequences, while decreasing the accuracy at the same time by adding false positives.

This is best seen in the drastic increase in homologous sequences within the DrosoSetupS1 when no synteny criterion is applied. Without an applied synteny criterion, nearly all neORFs in both setups exhibit homologous sequences in almost all target species. Applying synteny windows of different sizes influences the number significantly, suggesting that several of the identified homologous sequences detected with large windows could be false positives or the result of a bigger genomic reshuffling. For example, neORFs can be very small (Domazet-Loso and Tautz, 2003; Toll-Riera et al., 2009; Broeils et al., 2023) and sometimes be associated with transposable elements (Poretti et al., 2023; Lebherz et al., 2024a), in such a way that without synteny window, an ORF can be detected several times in a genome. Moreover, several studies have shown that *de novo* genes at early stages can undergo duplication (Grandchamp et al., 2023), leading them to be detected at different genomic locations. Additionally, little is known about duplication in non-genic regions (Bensasson et al., 2003; Xu et al., 2023), which may contribute to the high rate of detected homology across genomes.

Comparing the simple reciprocal BLAST hits with only the best reciprocal BLAST hits, also highlights the importance of a suited homology detection method. The percentage of syntenic homologous sequences of neORFs identified through simple reciprocal BLAST hits is higher in target species of the HumanSetupS1 than in populations of *Drosophila melanogaster* in the DrosoSetupS1. This finding is in contrast to the assumption that there should be a higher similarity between populations of the same species that between different species. However, comparing these numbers to the best reciprocal BLAST hits, we observe that the percentage of syntenic homologous sequences drops drastically in the HumanSetupS1, while the results remain almost unchanged in the DrosoSetupS1. This finding confirms that reciprocal BLAST hits recover members of larger gene families, while best reciprocal BLAST hits recover better the corresponding orthologous sequences (Hernández-Salmerón and Moreno-Hagelsieb, 2020; Moreno-Hagelsieb and Latimer, 2008). Since mammalian gene families are, on average, larger than *Drosophila* gene families (Hahn et al., 2007; Dornburg et al., 2022), this hypothesis could explain the observed patterns in the results between the HumanSetupS1 and DrosoSetupS1.

Our findings therefore emphasise the need for suited synteny methods to validate homologous sequences of neORFs, fitting the used input data and phylogenetic setup. Full genome alignments can provide a more comprehensive approach for synteny detection (Wacholder et al., 2023). However, whole genome alignments are not applicable for every phylogenetic setup and costly or difficult to get (therefore we go a middle way). **DESwoMAN** assesses synteny by using annotated genes as anchors for intergenic neORFS. However, alternative anchors become available for assessing synteny(Käther et al., 2025). Nevertheless, our results show that our method identifies a large proportion of syntenic homologous sequences well.

Regardless of the applied synteny method or window size or usage of best reciprocal BLAST hits, our results confirm that the phylogenetic distance is one of the most important factors for the detection and validation of neORFs and their corresponding homologous sequences. In the mouse, as a far-related species, for example, we could detect homologous sequences for only 18% of human neORFs even without a synteny criterion applied.

### Strategy 2: Tissue-specificity of neORFs in human

Another use case of **DESwoMAN** is to investigate the tissue-specificity of neORFs, which is implemented in an automatised way through Strategy 2. As a biological use case we utilise a different Human Setup (HumanSetupS2) with transcriptomes from five different tissues. We observe a universal pattern across all neORF categories (antisense, intronic, and intergenic) identifying the vast majority of them to be tissue-specific with a decreasing number of neORFs shared between an increasing number of tissues. This result suggests that transcription of new ORFs is a more complex event and therefore rare compared with the emergence of other coding features in an ORF. This result aligns with previous findings by Wacholder et al. (2023) or Grandchamp et al. (2024), who show that *de novo* transcript gain and loss is a highly dynamic process, with fast gain and loss processes. Understanding the gain, loss, and fixation of new transcription events presents a crucial challenge for a deeper understanding of neORF gain and fixation. **DESwoMAN** helps investigating the birth of neORFs independently of their fixation status. Therefore, our tool can help to advance our understanding of the earliest stages of *de novo* gene emergence and, with its transcriptomic focus, facilitate insights into the role of transcription in this process.

## Conclusion

Understanding how *de novo* genes emerge - from initially non-coding sequences to functional genes - is a critical yet still poorly understood aspect of genome evolution.

These genes often encode proteins with entirely novel functions, making them particularly relevant not only for understanding evolutionary processes, but also for applications in biotechnology and protein engineering. A deeper understanding of these early molecular events could illuminate the evolutionary origins of gene families, how life on earth emerged and evolved and support the rational design of novel proteins in synthetic biology.

Until now, research on *de novo* gene evolution has largely focused on genes that have already been fixed within a species, typically exhibiting stable transcription and complete coding features. However, the earliest stages, where non-coding sequences first gain transcriptional activity and gradually acquire coding potential, remain largely unexplored.

To address this gap, we developed **DESwoMAN**, a tool designed to detect and validate newly expressed open reading frames (neORFs) from transcriptome data and analyse the mutations involved in their emergence by comparing them with identified syntenic homologous sequences. To our knowledge, **DESwoMAN** is the first fully automated pipeline capable of performing these analyses based on transcripts, enabling systematic investigations into the initial steps of *de novo* gene formation.

**DESwoMAN** is highly customisable, allowing users to define analysis strategies and parameters according to specific research questions and dataset characteristics. In this study, we applied **DESwoMAN** to three distinct use cases, demonstrating its utility for investigating neORF evolution at the species, population and tissue-specific level and highlight biologically relevant factors and the impact of specific parameters.

**DESwoMAN** is intended to serve the scientific community as a valuable resource for gaining deeper insights into the early molecular mechanisms underlying *de novo* gene emergence and for supporting future work in evolutionary genomics and synthetic biology.

## Methods

### DESwoMAN Implementation and Parameters

**DESwoMAN** is implemented in Python 3. It requires three non-Python dependencies - BLAST (Altschul et al., 1990), gffread (Pertea and Pertea, 2020) and diamond (Buchfink et al., 2015) - for which reason we provide a docker image on DockerHub to run DESwoMAN directly with all dependencies installed. As input for **DESwoMAN** the user has to provide a set of query and target genomes and transcriptomes from different species, populations, or biological samples. Based on an input data set, **DESwoMAN** can be run with one of two different strategies (Fig. 4).

**Figure 4:**
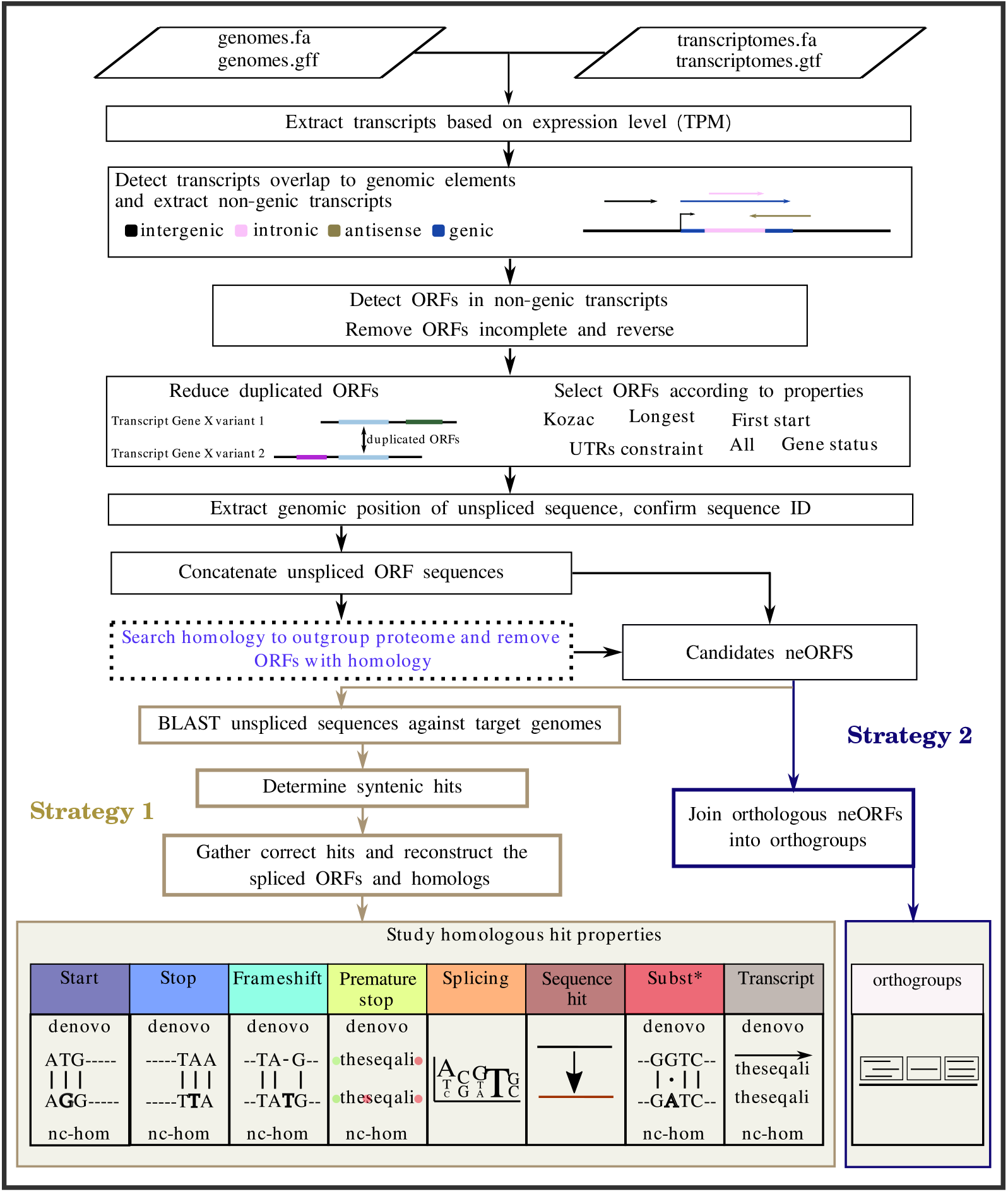
Flowchart of the DESwoMAN methodology.

The two strategies and their respective analysis steps are described in more detail below. A more detailed documentation of both strategies and all parameters of **DESwoMAN** can be found in the manual (Supp.Data.1), and in the readme of the github repository (https://github.com/AnnaGrBio/DESWOMAN).

### Strategy 1

The input data required for Strategy 1 is one query genome and a corresponding transcriptome. At least one closely-related target genome is mandatory, while corresponding target transcriptomes to any target genome are optional. Furthermore, a dataset of protein or DNA sequences is optional and recommended in case of homology search (Fig. 4).

With Strategy 1, neORFs are identified in a single query transcriptome. Different user-defined criteria can be applied for the extraction of candidate ORFs from the transcripts. Among all extracted ORFs, neORFs are retained if they lack similarity to genes from outgroup species. Additionally, neORFs are validated through syntenic non-genic homologous sequences in outgroup genomes. These syntenic non-coding counterparts are used to detect mutations in the neORFs (not) leading to a coding status. The following coding features of neORFs are inspected and reported by **DESwoMAN**: presence of a start codon, presence of a stop codon, frameshift mutation score based on the method of Wacholder et al. (2023), presence of a premature stop codon, number of substitutions, and presence of transcription if target transcriptomes are provided.

Several parameters and criteria of **DESwoMAN** can be user-defined based on the specific biological research question and input data. More details and recommendations regarding these options and their implementation are available in manual (Supp.Data.1).

### Strategy 2

The input data needed for Strategy 2 are at least two query transcriptomes assembled with reference-based algorithms (Raghavan et al., 2022; Kovaka et al., 2019) to a single reference genome. With Strategy 2, neORFs are identified in all query transcriptomes in contrast to only one as in Strategy 1. Strategy 2 groups neORFs from the multiple query transcriptomes into orthogroups. The orthogroups allow to study whether neORFs are specific to a transcriptome or rather expressed in several individuals, populations or conditions according to the selected transcriptomes. The identification of ORFs and the selection of neORFs based on homology is the same between Strategy 1 and 2.

As for Strategy 1, several mandatory and optional parameters and criteria of **DESwoMAN** can be user-defined based on the specific biological research question and input data. More details and recommendations regarding these options and their implementation are available in manual (Supp.Data.1).

### Use Cases: Human and Fruit Fly neORFs

To illustrate the use of **DESwoMAN**, we applied it to biological datasets with the two different strategies. Strategy 1 was applied to two setups: a Human Setup called “HumanSetupS1” with *Homo sapiens* as the query genome/transcriptome(s) and six mammalian target genomes/transcriptomes; and a Fruit Fly Setup “DrosoSetupS1” with *Drosophila melanogaster* as the query genome/transcriptome(s) and six other *D. melanogaster* samples from different geographical origins as target genomes/transcriptomes. Strategy 2 was applied to a new Human Setup “HumanSetupS2” with 5 transcriptomes from *Homo sapiens* from different tissues. All assembled transcriptomes and dataset generated can be found in https://zenodo.org/records/14936107.

### HumanSetupS1

For the HumanSetupS1, the genome of *Homo sapiens* (human): GRCh38.p14 from the NCBI RefSeq database (O’Leary et al., 2016) is used as a query genome.

Six target genomes and transcriptomes are used: As target genomes, the reference genomes of *Pan Paniscus* (Bonobo) : NCBI RefSeq assembly GCF_029289425.2, *Gorilla gorilla* (Gorilla) NCBI RefSeq assembly GCF_029281585.2, *Pan Troglodytes* (Chimpanzee) NCBI RefSeq assembly GCF_028858775.2, *Macaca mulatta* (Macaque) NCBI RefSeq assembly GCF_003339765.1, *Mus musculus* (Mouse) NCBI RefSeq assembly GCF_000001635.27 and *Pongo Pygmaeus* (Orangutan)

NCBI RefSeq assembly GCF_028885625.2 are used. For each genome, high-quality polyadeny-lated RNA-seq libraries of the brain (Brawand et al., 2011) from the NCBI Sequence Read Archive (Leinonen et al., 2010) are used as corresponding transcriptomes. *Homo sapiens*: brain (ID hsa br F 1 SRR306838); *Mus musculus*: brain (ID mmu br F 1 SRR306757); *Gorilla gorilla* brain (ID ggo br M 1 SRR306801); *Pongo pygmaeus* brain (ID ppy br F 1 SRR306791); *Macaca mulatta* brain (ID mml br F 1 SRR306777); *Pan troglodytes* brain (ID ptr br F 1 SRR306811); *Pan paniscus* brain (ID ppa br F 1 SRR306826).

All RNA-seq data are assembled using mapping-based assembly methods. Reads are trimmed of adapters and low-quality bases (quality scores < 15, minimum size kept: 36 nucleotides) using Trimmomatic (Bolger et al., 2014). Reads were then converted to FASTA format with seqtk (https://github.com/lh3/seqtk). The reads from each species were mapped to the corresponding reference genome.

All reference genomes are indexed with HISAT2 (2.2.1) using the “-build” module (Kim et al., 2019), and reads are mapped to the corresponding genomes using the HISAT2 with default parameters. The resulting SAM files are converted to BAM format, sorted, and indexed with SAMtools (1.13) (Li et al., 2009). The GTF annotation files of transcriptome assemblies are built with StingTie (1.3.4d) (Pertea et al., 2015). Conversion of transcriptomes to FASTA format is done with GffRead (Pertea and Pertea, 2020).

We built two different datasets for the homology search part performed by DESwoMAN: one containing protein sequences and one containing ncRNA sequences. Both datasets contain sequences of several non-mammalian outgroup species (Supp.Data.2.1).

### DrosoSetupS1

For the DrosoSetupS1, genomes and transcriptomes from seven samples of *Drosophila melanogaster* collected in different locations by the European Drosophila Population Genomics Consortium (FI: Finland, DK: Denmark, ES: Spain, SE: Sweden, UA: Ukraine, TR: Turkey, and ZI: Zambia) are used from Grandchamp et al. (2023). Details about the genome and transcriptome sequencing, assembly and mapping can be found in Grandchamp et al. (2023).

For the homology search, we built two different datasets with again protein and ncRNA sequences. Both datasets contain sequences of several outgroup species of *Drosophila melanogaster* (Supp.Data.2.2).

### HumanSetupS2

For the HumanSetupS2, five human RNA-seq libraries (Brawand et al., 2011) from different tissues are used as query transcriptomes: brain (ID hsa br F 1 SRR306838), cerebellum (ID hsa cb F 1 SRR306844), heart (ID hsa ht F 1 SRR306847), kidney (ID hsa kd F 1 SRR306851) and testis (ID has ts M 1 SRR306857). The same human reference genomes as for HumanSetupS1 is used to assemble the 5 transcriptomes. All RNA-seq data are assembled as explained in HumanSetupS1. All human reads were separately mapped to the human reference genome to generate 5 different transcriptomes.

We built two different datasets for the homology search performed by DESwoMAN: one containing protein sequences and one containing ncRNA sequences (Supp.Data.2.1). Closely related mammalian species were implemented into the 2 reference datasets from the HumanSetupS1 (together with the more distant non-mammalian outgroups). The list of species implemented can be found in (Supp.Data.2.3).

### Strategy applications for setups

First, Strategy 1 was run on HumanSetupS1 and DrosoSetupS1 with a synteny window of 3. Only intergenic transcripts were retained, with an expression threshold of 0.5 TPM. The ORFs selected in transcripts were the longest ORF. All simple reciprocal hits were used as an option to validate synteny. The stop codons were looked for in the first 50% of the sequences. The remaining parameters were default parameters.

Second, Strategy 1 was run with the same parameters, but all synteny windows were tested with simple BLAST hits and reciprocal BLAST hits for both setups.

Strategy 2 was run with the same parameters as Strategy 1 (synteny window of 3, simple BLAST hits), but additionally for intronic and antisense neORFs.

## Data access

**DESwoMAN** is available in https://github.com/AnnaGrBio/DESWOMAN. All result and output files, assembled mammalian transcriptomes, as well as Drosophila genomes and transcriptomes, can be found in https://zenodo.org/records/14936107 (version 1 and 2). A supplementary manual and document are in linked archived files.

## Acknowledgments

AG was supported by the Aix-Marseille University, by the Deutsche Forschungsgemeinschaft priority program “Genomic Basis of Evolutionary Innovations” (SPP 2349) BO 2544/20-1 to Erich Bornberg-Bauer, and by the Human Frontier Science Program Research Grant RGP004/2023 (doi.org/10.52044/HFSP.RGP0042023.pc.gr.168590) awarded to Erich Bornberg-Bauer, Anne-Ruxandra Carvunis and Christine Brun. ML was supported by the Deutsche Forschungsgemeinschaft priority program “The genomic basis of evolutionary innovations” (SPP2349; Project No. 503272152 awarded to John Parsch [PA 903/12-1] and Erich Bornberg-Bauer [BO 2544/20-1]). ED was funded by the Deutsche Forschungsgemeinschaft (DFG, German Research Foundation) – 503348080. We thank Erich Bornberg-Bauer and Christine Brun for sharing part of their funding to support this project.

## Author Contributions

Project management was conducted by AG. AG developed **DESwoMAN**. ML improved parts of the code and tested the program. ED edited code for python packaging and developed the docker container. AG wrote the first draft of the paper, which was revised by all authors who also contributed to the interpretation of the data. The authors declare no conflict of interest.

